# A Noninvasive Approach to Optogenetics Using Focused Ultrasound Blood Brain Barrier Disruption for the Delivery of Radioluminescent Particles

**DOI:** 10.1101/2020.08.20.248302

**Authors:** Megan Rich, Eric Zhang, Ashley Dickey, Haley Jones, Kelli Cannon, Yuriy Bandera, Stephen Foulger, Farah Lubin, Mark Bolding

## Abstract

Optogenetics, the genetic incorporation of light-sensitive proteins such as Channelrhodopsin-2 (ChR2) into target mammalian neurons, has enabled activation, silencing, and receptor subtype specific neuromodulation with high spatiotemporal resolution. However, the essential components of the ontogenetic system require invasive procedures with very few non-invasive alternatives preventing its use as a translational tool. The implantation of light emitting fibers deep within brain structures is both technically demanding and causes tissue scarring in target brain regions. To overcome these limitations, while maintaining the highly-tuned components of optogenetics we have developed a novel noninvasive alternative. Our approach replaces fibers with light-emitting radioluminescent particles (RLPs) that can be activated non-invasively with X-ray exposure. Here, we report successful noninvasive delivery of RLPs to target brain regions using MRI-guided focused ultrasound (FUS) blood brain barrier opening. In addition, FUS BBBO can be used to deliver viral vectors for light sensitive channel expression. Combined, these components can provide a completely non-invasive optogenetic system.

## Introduction

Originally discovered in algae, the family of opsin proteins includes multiple 7-transmembrane domain proteins (termed rhodopsin) that are activated by light and perform a range of cellular functions including ion channels and pumps [1]. In the field of optogenetics, neurons are genetically engineered to express opsins along with fluorescent reporters, which provides identification and control over a single neuron, or populations of neurons[2],[3]. Since this seminal work[4], [5] many microbial opsins have now been developed for *in vivo* and *ex vivo* applications including channelrhodopsins (e.g. ChR2) and halorhodopsins (e.g. eNpHR3.0) providing both excitatory and inhibitory neuronal control, respectively. However, the essential components of the optogenetic system require invasive procedures that limit its utility in translational research.

Focused Ultrasound (FUS) can penetrate the skull and in combination with commercial ultrasound contrast agents such as Definity, can safely and reversibly open the blood brain barrier (BBB) at any target brain location[6]–[12][13]. This allows the passage of circulating agents across the BBB only in brain regions targeted with FUS, providing spatially precise delivery on the order of millimeters. Furthermore, FUS BBB opening can be used in combination with MRI contrast agents to confirm the location of delivery [9], [14]–[16]. Recently, it has been shown by our group [17] and others [18]–[20] that focused ultrasound (FUS) BBB disruption can be used to provide noninvasive delivery of adeno associated virus (AAVs) including AAVs for the expression of ChR2 [20]. However, this study still required the implantation of an optic fiber cable and therefore, was not a completely noninvasive optogenetic system [20].

The hardware to deliver light deep within the brain of freely behaving animals has been limited to integrated fiber optics, lasers[21], [22], and light emitting diodes (LEDs)[23]. Surgical insertion into the brain of a fiber optic waveguide or an LED, which can be hundreds of microns in width, has several problems. First, significant mechanical damage to delicate brain tissue[21], [22] causes glial scarring at the light source and can decrease the effectiveness of light intensities leading to variability in channel activation. In addition, the light intensities required to activate the neurons with fiber optic delivery can result in local heating of the brain tissue, potentially leading to thermal ablation and/or unwanted physiological effects[24]. Furthermore, light is delivered only at the tip of the light source and is highly absorbed and scattered in brain tissue especially at shorter wavelengths[25] such as those needed to activate ChR2[26].

In order to overcome these limitations, we are working to replace fibers with novel light-emitting core-shell nanoparticles that can be activated noninvasively with X-ray exposure and can be strategically deposited such that they provide a more uniform distribution of light [27], [28]. Though further testing is still needed, these radioluminescent particle (RLPs) do not appear to interfere with neuronal functioning [29]. Additionally, external X-ray activation can penetrate all regions of the brain and can be homogeneously applied to major brain structures.

RLP technology provides a non-invasive alternative to implanted light sources used in current optogenetic systems however, the presence of the BBB which prevents almost all large molecules including nanoparticles similar in size to RLPs [7] from entering the brain parenchyma [30] would require invasive injection of particles into target brain regions. Here we investigate the use of microbubble-assisted FUS BBB opening for the location specific delivery of RLPs in order to provide an entirely noninvasive optogenetic system. Here, we report the successful delivery of RLPs to target brain regions using MRI-guided FUS BBB opening. This noninvasive alternative to current optogenetic procedures provides the optogenetic system with translational utility.

## Materials and Methods

### Radioluminescent Particle Configuration - YSO:Ce-BSA Nanoparticles

Silica nanoparticles (70 nm) were synthesized by a modified Stöber process with tetraethyl orthosilicate, ethanol, deionized (D.I.) water, and ammonia hydroxide (30% v/v) in a molar ratio of 1 : 88 : 56 : 7.01 : 1.30 respectively, mixed for 20 hrs, and washed several times with ethanol and water. Yttrium and cerium hydroxide were deposited onto the silica particle with a base catalyzed reaction in a 1:1 molar ratio of rare earth nitrate hexahydrate and silica in an aqueous solution. The optimal cerium nitrate hexahydrate concentration was determined to be 0.75% of the total rare earth reagent. The pH was adjusted to 9.2 with ammonia hydroxide and mixed for 24 hrs. The particles were annealed in air for 1 hr at 1000°C.

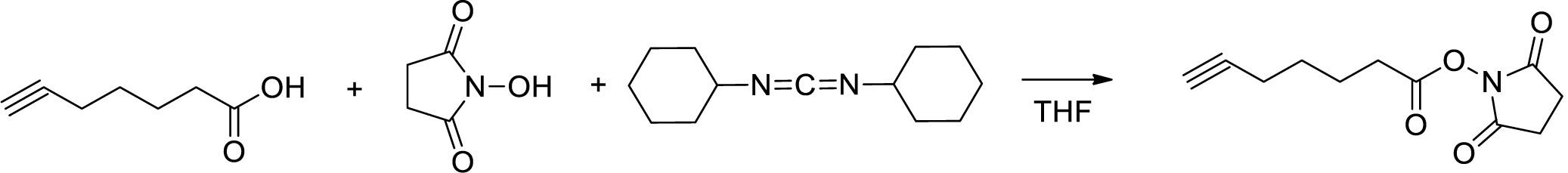

### Synthesis of 2,5-Dioxopyrrolidin-1-yl hept-6-ynoate (alkNHS)

An alkyne modified N-hydroxysuccinimide (NHS) derivative (alkNHS, 2,5-dioxopyrrolidin-1-yl hept-6-ynoate) was synthesized by dissolving hept-6-ynoic acid (1 g, 7.93 mmol) and N-hydroxysuccinimide (1 g, 8.72 mmol) in THF (25 mL). The solution was stirred and cooled with ice, then dicyclohexylcarbodiimide (1.8 g, 8.72 mmol) was added. The obtained mixture was stirred while cooling for 15 mins, at room temperature for 24 hrs and then filtered. The insoluble solid was washed with dichloromethane. The filtrate was evaporated under reduced pressure and the residue was dissolved in dichloromethane. The obtained solution was washed with water and the organic layer was separated and dried with Na_2_SO_4_, filtered, and evaporated under reduced pressure. The residue was dissolved in dichloromethane (10 mL) and cooled. Insoluble impurities were separated by filtration and the filtrate was evaporated to give a product with 90% purity by NMR. Yield 1.55 g, clear solid, mp = 55-56°C.

^1^H NMR (CDCl_3_) δ 1.65 (m, 2H, *J*=7.0 Hz), 1.87 (m, 2H, *J*=7.6 Hz), 1.97 (t, 1H, *J*=2.4 Hz), 2.25 (m, 2H, *J*=7.0 Hz, *J*=2.4 Hz), 2.65 (t, 2H, *J*=7.6 Hz), 2.84 (s, 4H).

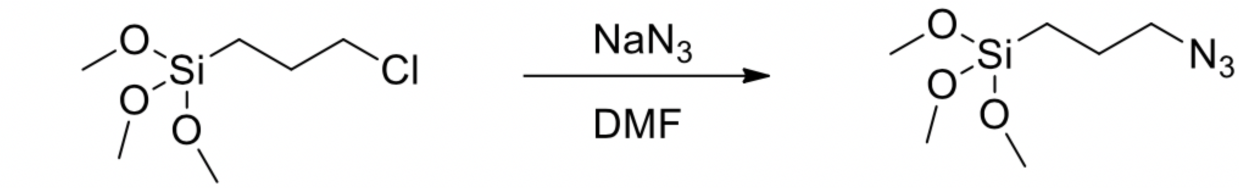

### Synthesis of (3-Azidopropyl)trimethoxysilane (azSil)

An azide modified silane linker (azSil, (3-azidopropyl)trimethoxysilane) was synthesized by a slightly modified, previously reported procedure[31]. (3-Chloropropyl)trimethoxysilane (1 g, 5.03 mmol) and sodium azide (0.684 g, 10.1 mmol) were mixed with dry dimethylformamide (3 mL) under nitrogen. The obtained mixture was stirred at 90°C for 1 hr. After cooling, the mixture was filtered. The filtrate solution contained 25% of product in DMF. ^1^H NMR (CDCl3) δ 0.68 (t, 2H, J=8.3 Hz), 1.70 (m, 2H, J=6.9 Hz), 3.25 (t, 2H, J=6.9 Hz), 3.56 (s, 9H).

Click chemistry was performed between the YSO:Ce particulates and bovine serum albumin (BSA) to passivate the particulates in the rodents, this chemistry is described briefly here. The core shell YSO:Ce particulates were modified with azSil to have azide functionality by dispersing 200 mg of the particles into 6 mL of dry dimethylformamide followed by the addition of 0.4 mL of water and 2 µL of azSil. 0.4 mL of ammonia hydroxide (28 % v/v) was added dropwise to initiate the reaction and mixed vigorously for 1 hr. In parallel, BSA was modified to have alkyne functionality (alkBSA) with standard N-hydroxysuccinimide coupling chemistry, where excess alkNHS was reacted with a 1 mM solution of BSA (500 mg in 7.52 mL 1x PBS) in a wrist-action shaker for 24 hrs in the dark. The resulting alkBSA was cleaned by dialysis in a Float-a-lyzer dialysis tube (8000-10000 MWCO) in water for 3 days with water changed frequently.

The azide modified particles and alkBSA were reacted in a copper(I)-catalyzed azide-alkyne cycloaddition (CuAAC) “click” reaction. In short, 150 mg of azide modified YSO:Ce particles were added to a J-KEM mini reactor tube equipped with a stir bar and dispersed in 2.350 mL D.I. water. Aqueous solutions of 4.52 mM copper sulfate (0.564 mg in 0.5 mL in D.I. water) and 11.3 mM sodium ascorbate (1.12 mg in 0.5 mL D.I. water) were added to the mini reactor tube followed by the addition of 75 mg of alkBSA in 1.650 mL 1x PBS. The reactor tube was placed in a J-KEM mini reactor vessel and allowed to react at 28°C for 24 hrs, under nitrogen, in the dark, and with stirring. After 24 hrs, the product was washed via centrifugation at 6500 rpm for 10 minutes two times with 1x PBS and 2 mL of ethylenediaminetetraacetic acid (EDTA) to remove the copper catalyst followed by two times with 1x PBS. The product was stored in 1x PBS at a concentration of 3.8 mg/mL in the fridge at 4°C.

### Radioluminescent Particle Configuration - SiO_2_/LPS:Ce,Tb,Eu-BSA-Fl Nanoparticles

A silica-lutetium pyrosilicate core-shell nanoparticle was synthesized using a high temperature multi-composite reactor, a technique that can overcome the temperature limitation nanoparticles can experience before sintering and is briefly described here. Silica nanoparticles were synthesized using the above Stöber method and kept in aqueous conditions. Silica nanoparticles (0.79g, 13mmol) were dispersed in 326 mL of water. Lutetium nitrate hexahydrate (1.4 g, 3.0 mmol), cerium nitrate hexahydrate (1.4 mg, 0.33 µmol), terbium nitrate hexahydrate (0.29 g, 0.65 mmol), and europium nitrate hexahydrate (0.29 g, 0.65 mmol) were then dissolved in the silica suspension. Sodium bicarbonate (1.1 g, 13 mmol) was dissolved in 285 mL of water and titrated into the silica/lanthanide suspension. The suspension was stirred for one hour and purified via centrifuge and D.I. water. Once dried the particulates were oxidized at 750°C for 30 mins. The SiO_2_/Lu_2_O_3_:Ce-Tb-Eu nanoparticles (1.45g) were redisperse in 218 mL of a 9:1 (v/v) methanol-water solution. 3-(trimethoxysilyl)propyl methacrylate (725 µL) and ammonia hydroxide (3.0 mL) was added to suspension and refluxed at 80°C for 4 hrs. The particulates were then dried using the above method and dried. The surface modified nanoparticles (1.4 g) were then redisperse in acetonitrile (228 mL) with divinylbenzene (1.4 mL) and the initiator azobisisobutyronitrile (98 mg). The suspension was purged with N_2_ and reacted at 55°C for 24 hrs. Once purified with methanol and dried, the SiO_2_/Lu_2_O_3_:Ce-Tb-Eu/pDVB nanoparticulates were carbonized at 1200°C for 16 hrs using a continuous flow of N_2._ The residual carbon species were combusted at 800°C for 1 hr with a continuous flow of air. This process was used to synthesize nonaggregate SiO_2_/LPS:Ce-Tb-Eu. AzSil was added to the SiO_2_/LPS:Ce-Tb-Eu nanoparticles using the same chemistry described previously. The SiO_2_/LPS:Ce-Tb-Eu particulates (1.04 g) were dispersed in 18.3 mL of methanol and 0.68 mL of water, followed by the addition of azSil (0.58 mL) and reflux for 2 hrs.

Click chemistry was performed between the azide modified SiO_2_/LPS:Ce-Tb-Eu nanoparticles and a fluorescein tagged bovine serum albumin (BSA) conjugate. BSA was modified with alkyne functionality using the same chemistry described previously.

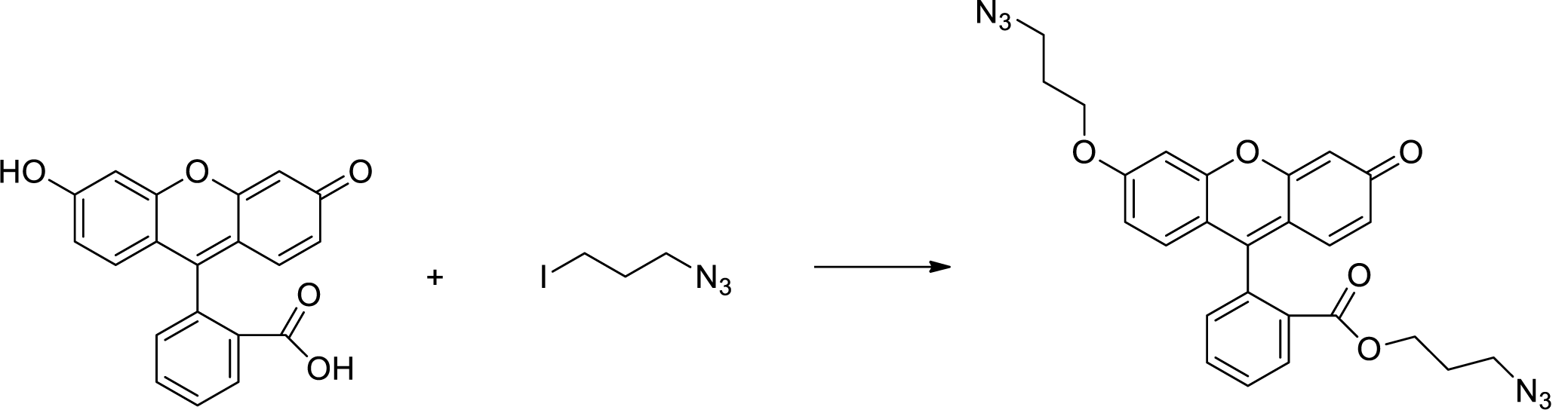

### Synthesis of 3-Azidopropyl 2-(6-(3-azidopropoxy)-3-oxo-3H-xanthen-9-yl)benzoate (azFl)

An azide modified fluorescein tag (azFl, 3-azidopropyl 2-(6-(3-azidopropoxy)-3-oxo-3H-xanthen-9-yl)benzoate) was synthesized by first dissolving fluorescein (0.4 g, 1.2 mmol) and 1-azido-3-iodopropane (0.56 g, 2.65 mmol) in dry dimethyl formamide (5ml) followed by the addition of potassium carbonate (0.5g, 3.6 mmol) under a nitrogen atmosphere. The mixture was stirred, heated to 80°C, and kept at this temperature for 4 hrs. After cooling, the mixture was extracted with dichloromethane and washed with water. The organic layer was separated and evaporated under vacuum. The obtained crude residue was mixed with water and the insoluble product was separated by decantation, dried under vacuum, and purified by column chromatography on silica. Solvent methanol:dichloromethane (1:20), Rf=0.2. Yield 0.55g (92%), orange crystals, m.p.=86-87°C.

^1^H NMR (CDCl3) δ 1.62 (m, 2H, J=6.2 Hz), 2.10 (m, 2H, J=6.2 Hz), 306 (t, 2H, J=6.2 Hz), 3.54 (t, 2H, J=6.2 Hz), 4.08 (t, 2H, J=6.2 Hz), 4.16 (t, 2H, J=6.2 Hz), 6.45 (d, 1H, J=2.4 Hz), 6.54 (d.d, 1H, J=2.4 Hz, J=9.6 Hz), 6.75 (d.d, 1H, J=2.4 Hz, J=8.9 Hz), 6.85(d, 1H, J=9.6 Hz), 6.89 (d, 1H, J=8.9 Hz), 6.97 (d, 1H, J=2.4 Hz), 7.31 (d.d, 1H, J=1.4 Hz, J=7.6 Hz), 7.65-7.78 (m, 2H, J=1.4 Hz, J=7.6 Hz), 8.25 (d.d, 1H, J=1.4 Hz, J=7.6 Hz).

The azide modified particles, alkBSA, and the azFl were reacted in a two-step copper(I)-catalyzed azide/alkyne cycloaddition (CuAAC) “click” reaction where, initially, the alkBSA and azFl were reacted to form a conjugate followed by the reaction of the conjugate with azide modified SiO_2_/LPS:Ce-Tb-Eu nanoparticles. Control over the reaction lies with controlling the functional groups that are available for clicking with the at a given time. In short, a 7.1 mM solution of azFl in tetrahydrofuran (THF) (26.55 mg in 7.5 mL THF) was added to a J-KEM mini reactor tube equipped with a stir bar followed by the addition of aqueous solutions of 4.39 mM copper sulfate (3.75 mg in 3.425 mL D.I. water) and 10.98 mM sodium ascorbate (7.45 mg in 3.425 mL D.I. water). The reactor tube was placed in the J-KEM mini reactor vessel and allowed to react for at 28°C for 1 hr under nitrogen, in the dark, and with stirring. After 1 hr, 1000 mg of the azide modified particles dispersed in 5 mL D.I. water was added to the reaction. The reaction was allowed to run for an additional 23 hrs in the same reaction conditions. After 23 hrs, the product was washed via centrifugation at 4000 rpm for 10 minutes two times with a solution of 1x PBS and 2 mL of EDTA to remove the copper catalyst, ten times with a solution of 12 mL THF and 18 mL 1x PBS until the supernatant was clear of dye by UV/Vis spectroscopy, followed by two times with D.I. water, and lastly once with 1x PBS. The final SiO_2_/LPS:Ce,Tb,Eu-BSA-FL product was stored in 1x PBS at a concentration of 85 mg/mL in the fridge at 4°C.

### MRI Procedure

For a detailed protocol on MRI and FUS procedures please refer to our published protocol [17]. Male Sprague Dawley rats weighing 250-300g were obtained from Charles River Laboratories and housed in accordance with UAB Institutional Animal Care and Use Committee (IACUC) guidelines. Rats had free access to water and rat chow, and were maintained on a 12:12 light:dark cycle. Animals were anesthetized with isoflurane (2 – 3%) and placed in a 3D printed stereotaxic frame, equipped with an MRI fiducial [17], [31]. Animals were maintained under isoflurane anesthesia throughout the MRI and FUS procedures. To confirm location and degree of BBB opening rats were MR imaged on a 9.4 T Bruker horizontal small-bore animal MRI scanner with a custom surface coil. T_1_- and T_2_-weighted images were collected prior to FUS procedure (prescan) and 15 minutes after. Image parameters were as follows, for T_1_- and T_2_-weighted axial images; width: 30 mm, height: 51.2 mm, depth: 3 mm, voxel size: 0.2x 0.2x 0.2 mm^3^, number of slices: 13; for T_1_-weighted coronal images; width: 30 mm, height: 30 mm, depth: 27 mm, voxel size: 0.2x 0.2x 1 mm^3^, number of slices: 27; for T_2_-weighted coronal images; width: 30 mm, height: 30 mm, depth: 27 mm, voxel size: 0.1x 0.1x 1 mm^3^, number of slices: 27. A pneumatic pillow sensor placed under the rat chest and connected through an ERT Control/Gating Module (SA Instruments) was used to monitor the rat’s respiratory rate throughout the imaging session. During the prescan, hippocampal target coordinates were measured in 3 planes from the MRI fiducial which would later be used for FUS targeting.

### FUS BBB Opening and RLP Delivery

The FUS BBB opening procedure was performed as described previously [17], [31]. Briefly, animals remained in the 3D printed stereotaxic frame and were transferred from the MRI to the ultrasound equipment. A tail vein catheter was then inserted and hair was shaved from the scalp and further removed with Nair. Using an XYZ positioning system (Velmex, Bloomfield, NY) driven by custom software written in LabVIEW (National Instruments, Austin TX), the ultrasound transducer was positioned to target coordinates and a water bath was coupled to the scalp with US gel and filled with degassed water. Then, the animals were intravenously injected with 1mL/kg of 3% Evans blue dye (Sigma Aldrich) and allowed to circulate for 5 minutes prior to FUS exposure. The animals were slowly infused with 30µL/kg of Definity (Lantheus Medical Imaging Inc., N. Billerica, MA.) while FUS was applied to either the left or right hippocampus while the other hippocampus was untreated for use as an internal control. A single element FUS transducer provided by FUS Instruments (Toronto, ON, Canada, 75mm aperture, 60mm focal length, full-width half-maximum intensity beam width: 1.14mm, full-width half-maximum intensity beam length: 5.99mm) was used. The FUS parameters were as follows: 0.30 MPa pressure in water, 1.1 MHz, 10 ms burst, 1 Hz burst repetition rate, 2-minute duration. The Definity infusion and FUS exposure were repeated a second time following a 5-minute gap allowing for the first Definity injection to clear from circulation (Bing et al. 2014). For RLP delivery, immediately following FUS, animals were IV injected with 15mg/kg of RLPs (SiO_2_/LPS:Ce,Tb,Eu-BSA-fluorescein, YSO:Ce-BSA, 70-80nm) in saline. All animals were also IV injected with 0.1 mL/ kg of Gadovist (Bayer Inc., Mississauga, Ontario, Canada) immediately after FUS for MRI confirmation of BBB opening.

### Brain Tissue Processing and Immunofluorescence

Animals were sacrificed for histological analysis at either 1 hour or 24 hours post RLP injection. Rats were perfused with 4% buffered formalin and brain tissue was immediately collected and frozen in Optimal Cutting Temperature compound (O.C.T.,Tissue-Tek, Sakura Finetek USA). Brain tissue was stored at -80°Cuntil cryostat sectioning. 10 µm frozen sections were collected and fixed with 4% buffered formalin, then rinsed with 1X PBS and coverslipped with either ProLong Gold mounting medium (Invitrogen, Carlsbad, CA) or Vectashield Mounting Medium with DAPI Hardset (Vector Laboratories, burlingame, CA). Neuronal marker NeuN staining protocol was performed as described previously [32]. Fluorescent microscopy imaging was performed using a Nikon 80i fluorescence microscope and processed using FIJI.

## Results

### FUS Mediated delivery of YSO:Ce-BSA

Successful FUS mediated BBB opening was confirmed 15 minutes following FUS application via enhanced Gadovist MRI contrast in the postscan image (Figure 1b) compared to the prescan image (Figure 1a). In order to determine if RLPs were able to enter the brain following FUS BBB opening, brain slices were examined under fluorescent microscopy. The initial particle configuration shown in Figure 1, YSO:Ce-BSA, are a blue-shifted variant and can be excited by UV exposure. Particles were apparent in the DAPI channel (Figure 1e,k) in the location of BBB opening which was evident by the EBD fluorescence (Figure 1d,j) indicating successful FUS-mediated delivery to the hippocampus.

**Figure 1.**
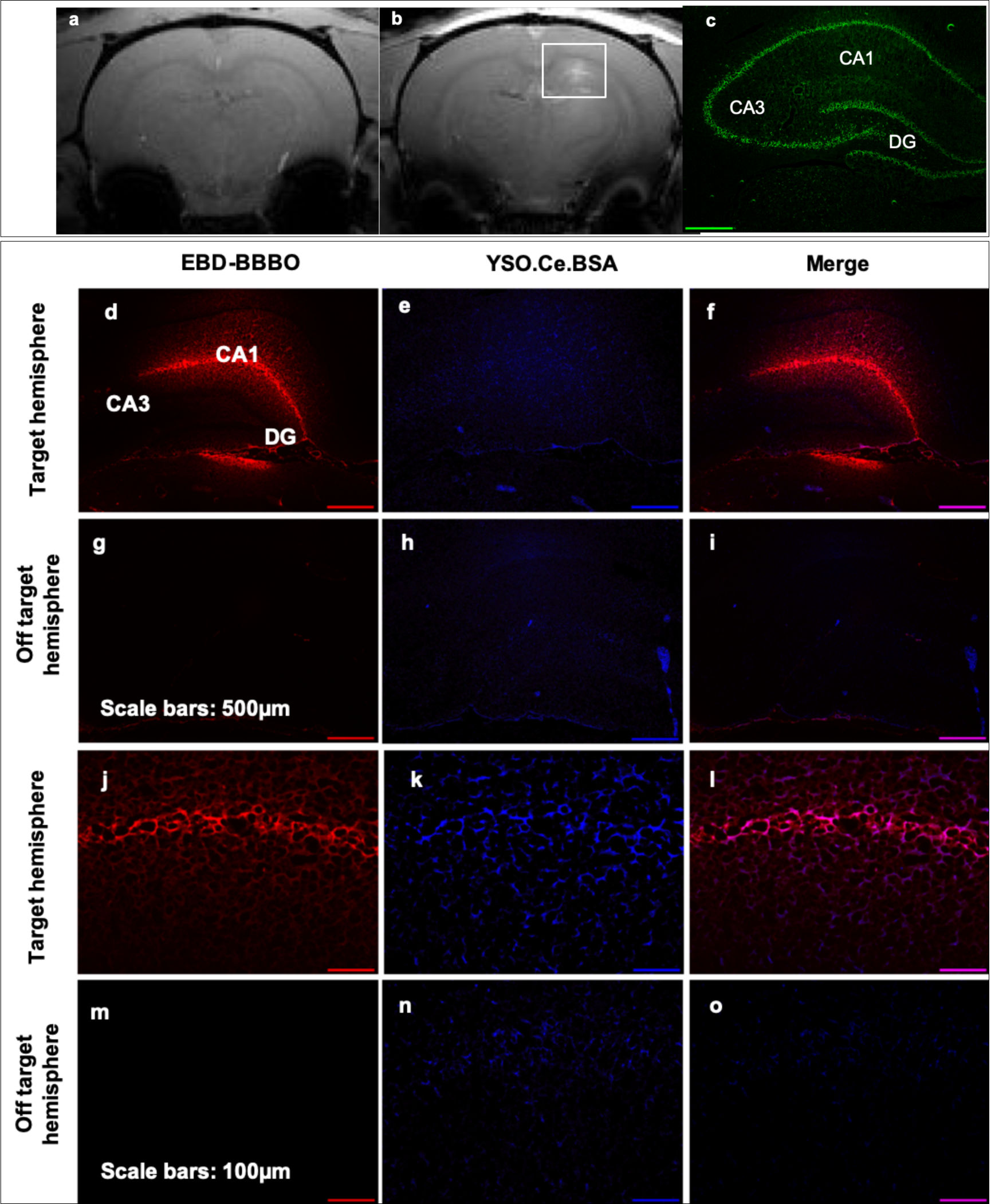
FUS-mediated delivery of YSO:Ce-BSA PLPs to the hippocampus. (a) MRI prescan (b) MRI postscan showing Gadovist MRI contrast in the FUS targeted hippocampus (white square). (c) anti-NeuN staining outlining hippocampal anatomy (CA1, CA3, DG). (d, j) FUS BBB opening further confirmed by Evans blue dye (EBD) in the target hemisphere. (e, k) Successful delivery of YSO:Ce-BSA RLPs was confirmed under UV exposure (blue) in the target hemisphere as compared to the off target hemisphere (h,n). Images in bottom row (j-o) are 5x the magnifications of top row (a-i).

### FUS Mediated delivery of SiO_2_/LPS:Ce,Tb,Eu-BSA-FL RLPs

Different RLP dopants produce different emission wavelengths and therefore can be used to activate different opsin proteins. For example, upon activation with x-ray, the Ce dopant produces a blue-shifted emission (460nm), making it a candidate for ChR2 activation, whereas the Ce-Tb-Eu dopant produces a redshifted emission (590nm) making it a candidate for ReaChR (Red-activatable variant of ChR2) or Chrimson opsin activation. Therefore, it is important to use the opsin protein and particle combination that will provide the most robust cellular response. The red-sifted Ce-Tb-Eu dopant particle variant provides a consistent emission output upon X-ray exposure (Figure 2). Furthermore, this variant can be used to excite the opsin ReaChR which has been shown to be more sensitive to some other opsin proteins [33]. In order to test whether these particular particles could be noninvasively delivered to target brain regions, animals underwent the FUS BBB opening procedure followed by IV injection with the SiO_2_/LPS:Ce,Tb,Eu-BSA particles that were tagged with fluorescein for later detection by fluorescent microscopy. Animals were then sacrificed at 1hr and 24hrs post injection for brain tissue collection. Our results show that the particles were present in the FUS targeted region (indicated by EBD expression) at both 1hr and 24hr time points (Figure 2). Note that at the 24hr time point the particles appear to have less expression compared to the 1hr time point suggesting some clearance may have occurred. Interestingly, at 24hrs the particles appear to tightly overlap with EBD which has been shown previously to clear from the brain progressively over 24hrs after FUS delivery via glial cell uptake via [34].

**Figure 2.**
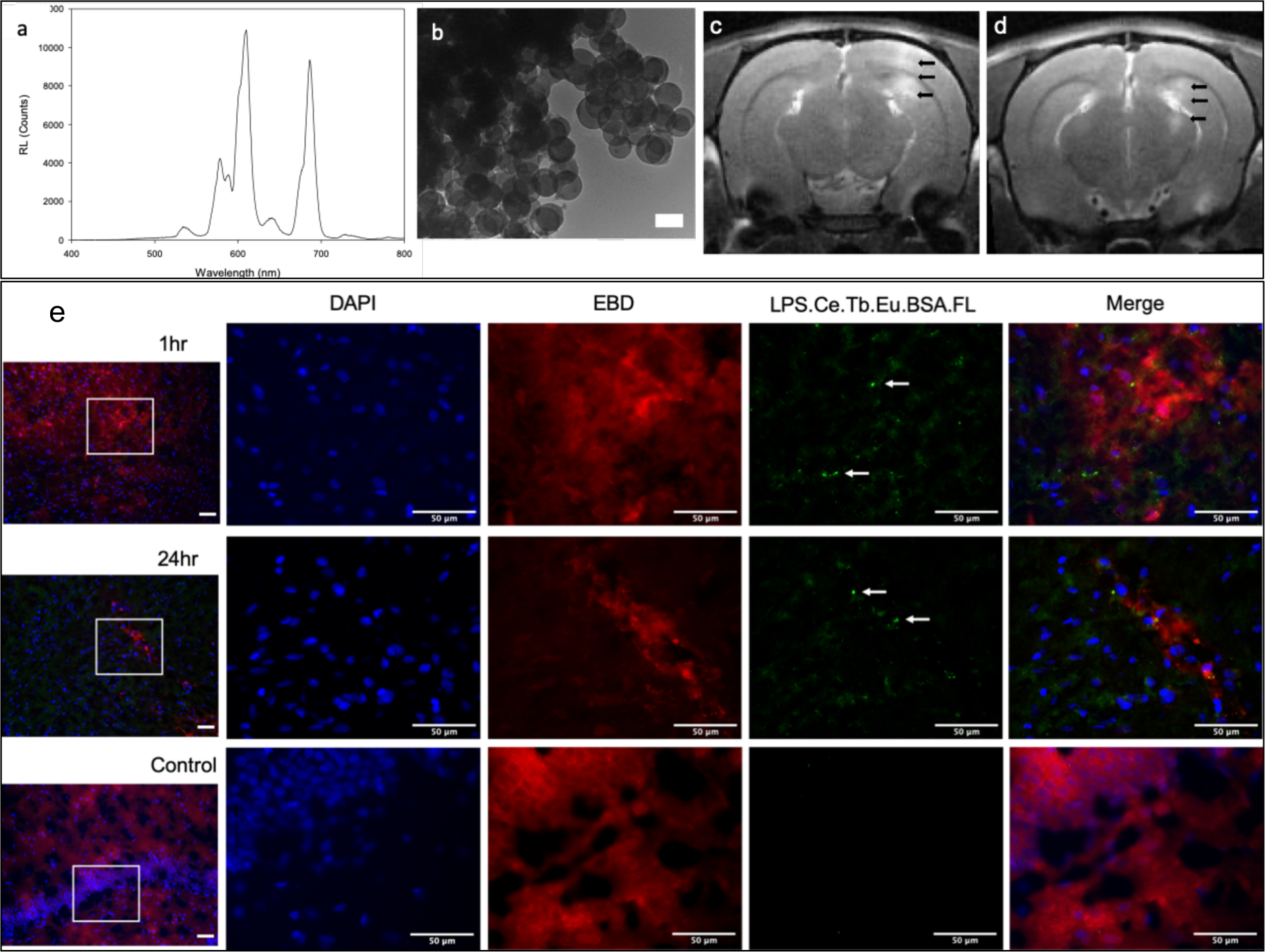
FUS mediated delivery of red-shifted RLPs. RLP emission wavelength upon x-ray activation of Ce,Tb,Eu particle variation (a). TEM image of SiO_2_/LPS:Ce,Tb,Eu RLPs, scale bar; 200nm (b). MR image showing BBB opening evident by Gadovist contrast (arrows) in animals injected with Red-shifted RLPs and sacrificed at 1hr (c) and 24hr (d). Histology showing RLPs were detected in the FUS targeted region at 1hr (top row) and 24hr (middle row) following delivery (e). Bottom row shows control tissue 1hr post FUS procedure without RLP injection. Left image inset represents the selected area of magnification for adjacent images. From left to right; (Blue) DAPI showing overall cellular morphology, (Red) Evans Blue Dye (EBD, red) showing region of BBB opening, (Green) SiO_2_/LPS:Ce,Tb,Eu. RLPs coated with bovine serum albumin (BSA) and tagged with fluorescein (Fl), (Merge) note RLPs (green) overlapping with EBD (red) in region of BBB opening at 1hr and at location of EBD clearance at 24hrs. All scale bars; 50 µm.

## Discussion

Optogenetics is a technique that is widely used in basic neuroscience studies due to its ability to produce cell-type specific activation, inhibition, or modification to neuronal activity with temporal precision on the order of milliseconds [4]. However, optogenetics currently requires invasive intracranial surgical procedures limiting its consideration for clinically translational applications. We investigated a novel optogenetic method that does not rely on any invasive injections or implants. We employ MRI-guided FUS BBB opening for the delivery of RLPs that can emit light upon exposure to X-ray energy. FUS can open the BBB in target locations on the order of millimeters, enabling entry of RLPs and viral vectors to specific brain regions including deep brain structures. We have shown previously that we were able to achieve location specific protein expression following FUS-mediated delivery of AAV9-hsyn-GFP [17]. And other groups have shown successful expression of opsin proteins following FUS mediated delivery of AAV9-mcherry-ChR2 [20]. However, in order to demonstrate that this noninvasive delivery method produced working opsin proteins Wang et al. needed to implant an optical fiber for light activation. RLPs provide an alternative to implanted optic fibers and light emitting diodes (LEDs) and have been shown to be nontoxic to neurons [27], [28]. Here, we confirm that we are able to noninvasively deliver RLPs to the rat hippocampus using FUS BBB opening (Figure 1). Now that we have confirmed that both RLPs and viral vectors can be delivered noninvasively, future studies can deliver AAVs with FUS for the expression of channelrhodopsins like ChR2 or ReaChR, followed by particle delivery and x-ray exposure to complete this noninvasive optogenetics platform. Prior to behavior studies however, additional studies should be conducted either in electrophysiological brain slice recordings or with histology to confirm x-ray activation of FUS-delivered RLPs, provides adequate light emission to activate FUS-delivered viral opsin protein constructs.

Many different opsin proteins have been developed and optimized for in vivo use, providing a diverse toolkit that enables a wider-range of neuronal manipulations [35], [36]. Each opsin protein is activatable by particular wavelength of light some providing better tissue penetrance and distribution than others [35]. Here, we investigate FUS-mediated delivery of two different RLPs, one producing a blue-shifted (470nm) emission and one producing a red-shifted emission (590nm). Thereby, matching the excitation peaks of the widely used cation channel ChR2 and ReaChR, the red-activatable variant of CrChR2, respectively. Although, hyperpolarizing variants of Channelrhodopsins do exist, they are mostly depolarizing cation channels. Recently however, additional RLPs have been developed with emission peaks around 540-570nm, matching the excitation spectrum of a commonly used inhibitory opsin protein, halorhodopsin [37]. It is important to note that the in vivo efficiency of this noninvasive system will depend on the particle distribution, intensity and tissue absorption of light output as well as the sensitivity of the opsin protein variant. Therefore, prior to intensive in vivo application, it is important to test the feasibility of this system with multiple variations of particles and proteins ideally with *ex-vivo* recordings.

## Conclusions

Here, we report successful noninvasive delivery of two different RLP variants to the hippocampus using MRI-guided focused ultrasound (FUS) blood brain barrier opening. In addition, FUS BBBO has been shown previously to provide successful delivery of viral vectors for light sensitive channel expression. Combined, these components can provide a completely non-invasive optogenetic system

## Acknowledgements

The authors thank the National Science Foundation (NSF) Grant # OIA-1632881 for financial support. In addition, this research was supported in part by the Civitan International Research Center, Birmingham, AL. Rich M. was supported by the Alabama EPSCoR Graduate Research Scholars Award. The authors gratefully acknowledge the use of the services and facilities of the UAB Comprehensive Cancer Center’s Preclinical Imaging Shared Facility Grant [NIH P30 CA013148] and the Vision Sciences Research Center core [NIH P30 EY003039].

